# Cortico-genetic mapping links individual brain maturity in youths to cognitive and psychiatric traits

**DOI:** 10.1101/487173

**Authors:** Tobias Kaufmann, Dennis van der Meer, Dag Alnæs, Oleksandr Frei, Olav B. Smeland, Ole A. Andreassen, Lars T. Westlye

## Abstract

Neurodevelopmental trajectories are shaped by interactions between coordinated biological processes and individual experiences throughout ontogeny, yet the specific genetic and environmental impact on brain development is enigmatic. Here, we map the genetic architectures of cognitive traits and psychiatric disorders onto the brain, show that such canonical genetic maps are associated with individual normative patterns in youths, and provide evidence that trauma exposure and parental education may alter this relationship.

---

Psychiatric, cognitive and brain imaging traits are highly heritable and polygenic^1–13^. Individual genetic architecture contributes to individual differences in neurodevelopmental trajectories and subsequent scaffolding and maintenance of brain structure and function throughout ontogeny. However, the links between the genetic and neural configurations and how their interplay shapes individual differences in cognitive function and mental health remain poorly understood. This knowledge gap has nurtured a debate on the extent of environmental influence and genetic constraints on brain development. Here, we provide a comprehensive neuroanatomical mapping of the genetic architecture of various cognitive traits and psychiatric disorders using brain imaging and genetic data in a large population based sample (*UK Biobank*^*14*^), and link the resulting canonical genetic maps to individual patterns of brain maturity in the *Philadelphia Neurodevelopmental Cohort*^*15*^.

We accessed data from the *UK Biobank*^*14*^ and used *Freesurfer*^16^ for cortical reconstruction based on T1-weighted magnetic resonance images obtained from 16,612 healthy individuals with European ancestry aged 40 to 70 years (mean: 55.8 years, sd: 7.5 years, 52.1% females). We computed surface maps for cortical thickness and area, registered to *fsaverage4* space (2,562 vertices), smoothed using a kernel with full width at half maximum of 15 mm. Next, we performed a genome-wide association study (GWAS) for every vertex using *PLINK*^*17*^, linking single-nucleotide-polymorphism (SNP) data with a given vertex’s thickness and area, respectively. Each GWAS accounted for effects of age, age^2^, sex, scanning site and the first four genetic principal components^18^ to account for population stratification.

We first estimated SNP-heritability of cortical morphology using *LD Score regression*^19^ for each vertex (**Suppl. Fig 1**). The spatial correlation between thickness and area heritability maps was moderate (r=0.28, p_*perm*_=0.0004; **Suppl. Fig 1b**), and surface area was significantly more heritable than thickness (**Suppl. Fig 1a**). Both measures showed regional differences, with high heritability of thickness in the postcentral gyrus and Heschl’s gyrus (**Suppl. Fig 1c**), and of surface area in the lingual gyrus and the temporal lobe (**Suppl. Fig 1d**). These results from vertex-wise SNP-based analysis largely confirmed earlier reports from twin studies^9–13^ and from a region-wise SNP-based analysis^20^, supporting the feasibility of our vertex-wise GWAS approach.

**Fig. 1:**
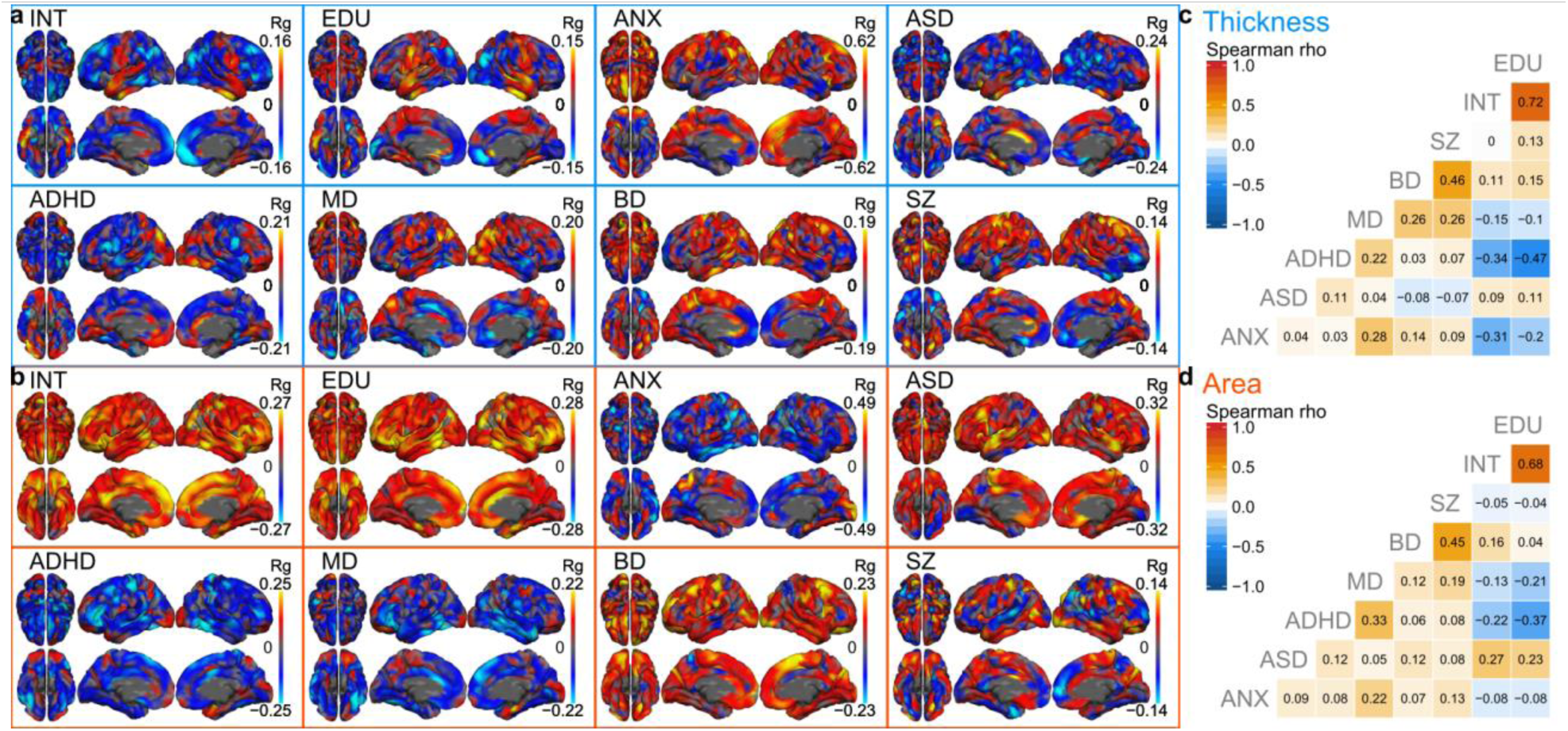
Cortico-genetic maps reflecting the vertex-wise genetic correlations of cortical morphology with cognition and psychiatric disorders. Genetic correlations (Rg) per phenotype for thickness (a) and for area (b). The maximum of the scales was individually adjusted to display the 97.5 percentile across all vertices. Corresponding p-values are depicted in **Suppl. Fig. 3.** (c-d) Pairwise Spearman correlations of the cortico-genetic maps from (a-b). Corresponding p-values from permutation testing are depicted in **Suppl. Fig. 4**.

We next combined our GWAS results with publicly available summary statistics to compute vertex-wise genetic correlations between brain morphology and cognitive traits and psychiatric disorders. Summary statistics for cognitive phenotypes were obtained from GWAS on intelligence^1^ (INT) and educational attainment^2^ (EDU), excluding *23andMe* data. For psychiatric disorders we used summary statistics from analyses on anxiety^3^ (ANX), autism spectrum disorder^4^ (ASD), attention-deficit-hyperactivity disorder^5^ (ADHD), major depression^6^ (MD, excluding *23andMe* data), bipolar disorder^7^ (BP) and schizophrenia^8^ (SZ). Using *LD Score regression*^19^, for each vertex we estimated the genetic correlation between each of the phenotypes and thickness and area, respectively. To reduce noise, vertices with a heritability estimate of less than 1.96 times its standard error were excluded from the analysis, in addition to excluding the medial wall, yielding a total of 4550 and 4498 vertices for thickness and area, respectively.

**Fig. 1a-b** depict the resulting cortical maps of vertex-wise genetic correlations – hereafter referred to as *cortico-genetic maps.* Each map reflects the overlap between the genetic architectures of cortical morphology and the given phenotype. For example, area in the right superior frontal gyrus was positively genetically associated with INT and negatively with MD, whereas thickness in this area was positively associated with ANX. **Fig. 1c-d** illustrate the correlation between these cortico-genetic maps, largely in line with the published genetic relationship between the phenotypes (**Suppl. Fig. 2** for comparison). Importantly, the cortico-genetic maps were derived from brain imaging data of healthy individuals, thereby reducing the impact of confounding factors such as comorbid disorders or medication, as might be observed for case-control brain imaging maps.

**Fig. 2:**
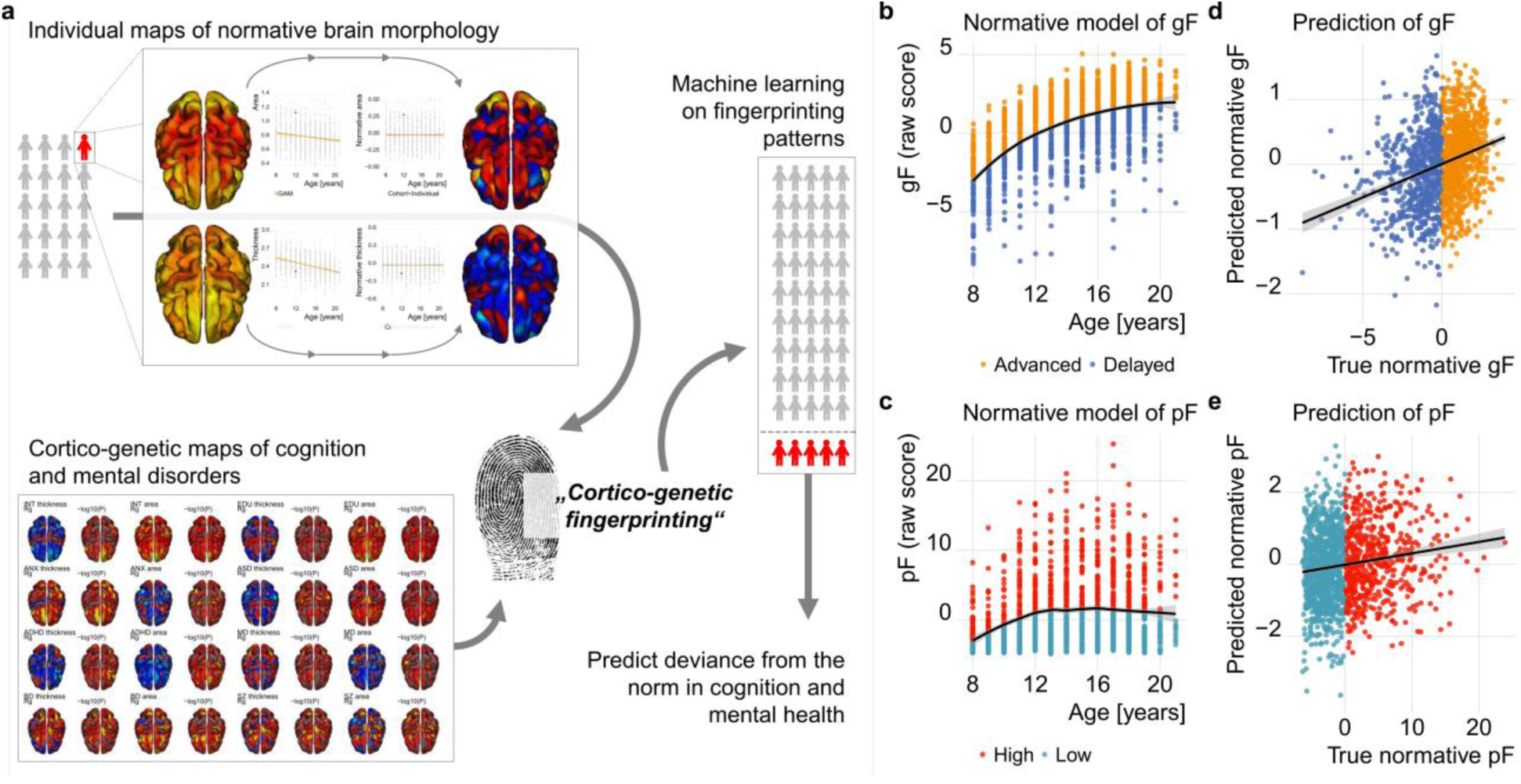
Cortico-genetic fingerprinting yields significant statistical predictions of normative general cognitive (gF) and general psychopathology (pF) factors in the developing brain. (a) Illustration of the cortico-genetic fingerprinting approach. (b) Association between general cognition factor (gF) and age. The black line indicates the model fit to remove the age effect, yielding normative estimates of gF. (c) Same as (b) but for general psychopathology (pF). (d-e) Significant prediction of normative gF (d) and normative pF (e) using cortico-genetic fingerprinting and machine learning. For corresponding permutation tests see **Suppl. Fig. 5**.

Considering the strong evidence of a neurodevelopmental component in the etiology of many psychiatric disorders^21,22^ and the large amount of maturational brain changes related to individual adaptation and learning^23^, we hypothesized that brain regions associated with the genetic architecture of psychiatric and cognitive traits in healthy adults are sensitive to normative deviations during childhood and adolescence. To this end, we tested if the similarity between an individual’s map of brain maturity and the respective cortico-genetic maps allowed us to statistically predict individual deviations from the developmental norm in the given trait in the *Philadelphia Neurodevelopmental Cohort*^*15*^. Following cortical reconstruction^16^, we excluded data due to insufficient quality after manual screening (n=60) and significant or major medical conditions (n=73), yielding a total sample of 1467 individuals aged 8 to 21 years (mean: 14.14 years, sd: 3.51 years, 52.9% females).

**Fig. 2a** details the approach. First, we modeled normative trajectories of brain development in each vertex using generalized additive models^24^ and removed the statistical relationship with age and sex from all data sets. For each individual, this yielded one normative map for cortical thickness and one for cortical area, where the measures at each vertex reflect its respective deviation from the age- and sex-matched sample norm. Next, we assessed the similarity of each individual’s normative thickness and area maps to each of the cortico-genetic maps from **Fig. 1a-b.** Following the *connectome fingerprinting* approach^25–27^, we hereafter refer to this as *cortico-genetic fingerprinting* outlining that the cortico-genetic maps are used as ‘*fingerprints*’ of cognitive traits and psychiatric disorders that individual patterns of normative development are compared to. Since the direction of effects in studies of brain morphometry may depend on age^28^, we also fingerprinted using the unsigned statistical maps. For example, whereas SZ is associated with widespread reduced cortical thickness^29^, the cortico-genetic thickness maps for SZ in **Fig. 1a** show positive genetic correlations in the precentral cortex. Speculatively, this may partly reflect a survivor bias as these maps were generated using data from healthy individuals aged 40 and above with very low life-time probability of a diagnosis. Therefore, in some cases the unsigned effect sizes might be needed to overcome the bias and to obtain predictive value in a neurodevelopmental sample. We thus fingerprinted using both signed genetic correlation maps (**Fig. 1a-b**) and unsigned maps of -log_10_ transformed p-values (**Suppl. Fig. 3**), yielding a total of eight (phenotypes) by three (area and thickness, and both concatenated) by two (genetic correlation, -log_10_ transformed p-values) fingerprinting correlations per individual (48 in total).

To assess the predictive utility of these fingerprints, we used machine-learning to predict deviations from the developmental norm in cognitive performance and mental health. Using principal component analysis on clinical and cognitive data, we derived a general cognitive score (gF) and a general psychopathology score (pF)^30^ and first removed effects of age using locally weighted regression (**Fig 2b-c**), yielding normative estimates of gF and pF. In a 10-fold cross-validation framework, we trained a linear machine learning model on 90% of the data to predict normative gF and normative pF using the 48 correlation estimates – the *connectome fingerprinting strength* - as features, and iteratively testing on the 10% held-out data. **Fig. 2d-e** illustrates that it was possible to statistically predict normative gF and normative pF using cortico-genetic fingerprinting (both p_perm_<.0001, **Suppl. Fig. 5**). In models accounting for age and sex, the associations between true and predicted normative values were highly significant both for gF (r_partial_=0.29; t=11.71, p=2e-16) and pF (r_partial_=0.14; t=5.51, p=4e-8). The machine learning model weights revealed that a range of traits contributed to each prediction (**Suppl. Fig. 6)**. These results jointly suggest that individual regional deviations from the norm in youths emerge in those cortical areas that are most strongly associated with the genetic architecture of the respective phenotypes, and that the extent of overlap with those patterns relates to individual differences in cognitive performance and – to a lesser degree - mental health.

The observed cortico-genetic overlap with individual estimates of brain maturity raises the question whether and to which degree this relationship is altered by experience. We used parental education as a proxy for the socioeconomic environment and the first component from a principle component analysis on trauma questionnaires as a proxy for major negative life events (**Suppl. Fig 7**). Next, we tested for linear associations between these factors and individual cortico-genetic fingerprinting strength. As depicted in **Fig. 3a**, parental education was positively associated with fingerprinting strengths on SZ (area), INT (thickness), EDU (thickness and area), and BD (area), as well as negatively associated with those on ASD (area) and ANX (thickness and area), each model accounting for age, sex and gF. Trauma exposure was positively associated with fingerprinting strengths on ANX (thickness), accounting for age, sex and a pF that excluded trauma items (**Fig. 3b)**. In other words, the overlap between an individual’s normative cortical morphology and the cortico-genetics maps from **Fig. 1** varied as a function of experience. With caution and under the restriction that these are probabilistic, not deterministic associations, these results indicate that individual developmental patterns are more ANX-like following trauma exposure, more ANX- and ASD-like in individuals from low-educated social environments and more SZ-, BD-, INT- and EDU-like in individuals from high-educated social environments.

**Fig. 3:**
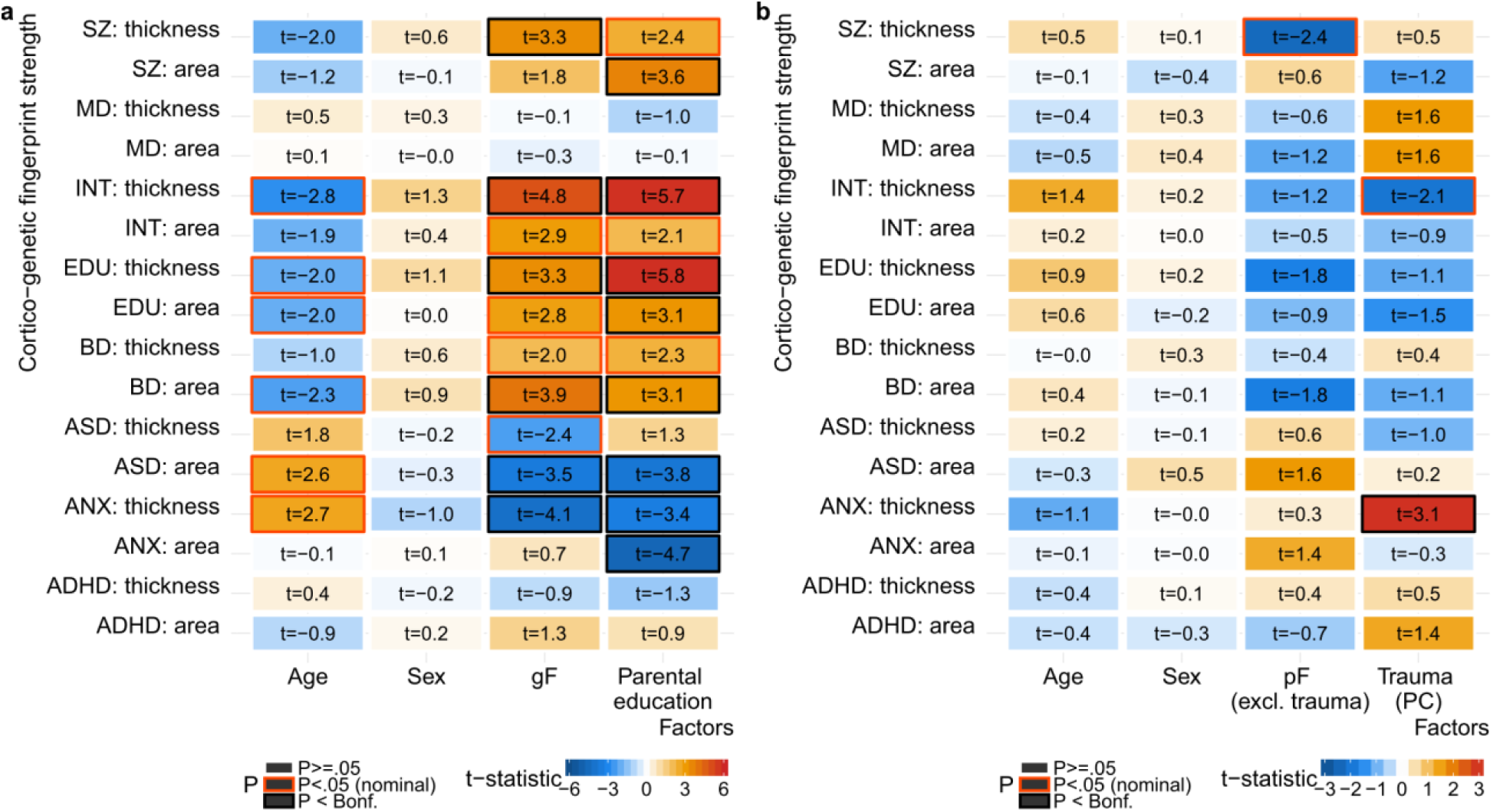
Association between individual cortico-genetic fingerprinting strength and proxies for socioeconomic environment and life events. *Cortico-genetic fingerprinting strength* refers to the correlation strength of each individual’s normative maps with the cortico-genetic maps from **Fig. 1**. For each *fingerprint*, we computed a linear model assessing (a) the association with parental education, accounting for age, sex and gF, and (b) the association with trauma, accounting for age, sex and pF (with trauma excluded from the pF). **Suppl. Fig. 8** displays permutation test results from 10,000 permutations. Significant effects at Bonferroni level (accounting for 16 maps, p=0.003125) for each factor are marked with a black box.

Taken together, our analysis reveals an intriguing impact of genetic architecture on brain development by illustrating that the similarity between individual patterns of brain maturity and the neurogenetics of cognition and psychopathology is informative for individual normative deviations in cognitive performance and mental health. Nevertheless, despite statistical significance our predictions only explained a proportion of the variance in the data, indicating that other factors have a large part in explaining individual trajectories. Indeed, we identified two environmental factors – proxies of the socioeconomic environment and adverse life events - as significant factors explaining variance in the individual fingerprints. Of note, these factors have substantial genetic components themselves, and future research needs to address to what extent the observed associations with environmental factors can be explained by common genetics. Apart from its utility in pinpointing deviations from the norm in the developing human brain, our cortico-genetic approach may contribute towards the delineation of genetic and environmental factors influencing individual trajectories during sensitive neurodevelopmental phases.

## Author contributions

T.K. and L.T.W conceived the study; T.K. and D.v.d.M. pre-processed the data. T.K. performed all analyses, with contributions from D.v.d.M and O.F. and with conceptual input from D.A. and L.T.W.; All authors contributed to interpretation of results; T.K. drafted the manuscript and all authors contributed to and approved the final manuscript.

## Acknowledgements

The authors were funded by the Research Council of Norway (276082, 213837, 223273, 204966/F20, 229129, 249795/F20, 248778), the South-Eastern Norway Regional Health Authority (2013-123, 2014-097, 2015-073, 2016-064) and Stiftelsen Kristian Gerhard Jebsen. This research has been conducted using the UK Biobank Resource under Application Number 27412 and the PNC under Application Number 8642. Support for the collection of the PNC data sets was provided by grant RC2MH089983 awarded to Raquel Gur and RC2MH089924 awarded to Hakon Hakonarson, and all subjects were recruited through the Center for Applied Genomics at The Children’s Hospital in Philadelphia.

## Online methods

### Samples and exclusion criteria

UK Biobank: The UK Biobank is a publicly available resource of imaging, genetics and phenotypic data from an ongoing large-scale cohort study^*14*^. All study procedures were approved by appropriate ethics committees and all study participants gave electronic signed consent. We obtained access with permission no. 27412. No statistical methods were used to pre-determine sample sizes as we analyzed all available data from the 20,000-subject release of brain imaging and corresponding phenotypic and genetics data. Individuals with Caucasian ancestry were identified by the UK Biobank study team using clustering analysis^18^ and we followed their selection of individuals in our study. After exclusion of data from individuals with a diagnosed brain disorder or data of insufficient quality (see *pre-processing and quality control*), this yielded a total of 16,612 healthy individuals with Caucasian ancestry. The age range was 40 to 70 years (mean: 55.8 years, sd: 7.5 years, 52.1% females).

PNC: The Philadelphia Neurodevelopmental Cohort is a publicly available resource of clinical, cognitive, genetic and neuroimaging data from children and adolescents^15,31^. Prior to data collection, all study procedures were approved by the institutional review boards of the University of Pennsylvania and the Children’s Hospital of Philadelphia, and all participants gave written informed consent. We obtained access with permission no. 8642. No statistical methods were used to predetermine sample sizes as we used all available data, except data with insufficient quality after manual screening (n=60) and data from individuals with significant or major medical conditions (n=73). The final sample comprised 1467 individuals aged 8 to 21 years (mean: 14.14 years, sd: 3.51 years, 52.9% females).

### Image pre-processing and quality control

T1-weighted magnetic resonance images for UK Biobank (MPRAGE, TR 2000 ms, TE 2.01 ms, matrix 208×256×256, resolution 1×1×1 mm) and PNC (MPRAGE, TR 1810 ms, TE 3.51 ms, matrix 192×256×160, resolution 0.9×0.9×1mm) was processed using Freesurfer 5.3^16^ (recon-all). In the case of UK Biobank, where manual quality control of 16,612 images was not feasible, we excluded outliers based on global cortical measures. We regressed age, age^2^, sex and scanning site from white surface area and mean cortical thickness of each hemisphere and identified outliers above or below 4 standard deviations of the full population, excluding N=22 individuals. In the case of PNC, we screened all reconstructed images manually, excluding data from N=60 children and adolescents that did not adhere to highest data quality standards. Next, for both UK Biobank and PNC, we resampled the surfaces to *fsaverage4* space (2,562 vertices), smoothed using a kernel with full width of half maximum of 15 mm.

### Vertex-wise genetic analysis in UK Biobank data

Standard quality control procedures were applied to the UK Biobank v3 imputed genetic data, including removal of SNPs with an imputation quality score below 0.5, with a minor allele frequency less than .05, missing in more than 5% of individuals, and failing the Hardy Weinberg equilibrium tests at a p<1×10^-6^. Genetic principal components were retrieved from the UK Biobank repository and we regressed the first four genetic components, age, age^2^, sex and scanning site from vertex-wise thickness and area maps. Next, we ran one genome-wide association analysis (GWAS) per vertex using *PLINK v1.9*^17^, and removed the MHC region from each resulting summary statistic. Using *LD Score regression*^19^, we estimated narrow sense heritability. The significance of the correlation between heritability maps of thickness and area was assessed using spin-rotation based permutation testing, which applies random rotations to spherical representations of the cortical surface to generate a null distribution^32^. Next, we used cross-trait *LD Score regression*^19,33^ to calculate correlations of our vertex-wise GWAS summary statistics with publicly available summary statistics on intelligence^1^ (INT), educational attainment^2^ (EDU, excluding *23andMe* data), anxiety^3^ (ANX, the case-control GWAS), autism spectrum disorder^4^ (ASD), attention-deficit-hyperactivity disorder^5^ (ADHD), major depression^6^ (MD, excluding *23andMe* data), bipolar disorder^7^ (BP) and schizophrenia^8^ (SZ). Significance of the correlations between each pair of the resulting cortico-genetic maps was again assessed using spin-rotation based permutation testing^32^ in addition to correcting the permuted p-values for the number of total correlations (28, Bonferroni correction).

### Cortico-genetic ‘fingerprinting’ in PNC data

We utilized generalized additive models^24^ to remove the statistical relationship with age and sex from vertex-wise thickness and area data, yielding one normative thickness and one normative area map per individual in PNC. Each of these individual subject maps was transformed into a one-column vector and correlated against similar vectors of the cortico-genetic maps of cognition and psychiatric disorders using Spearman correlations. We refer to this approach – in line with the *connectome fingerprinting* literature^25–27^ – as *cortico-genetic fingerprinting*. We fingerprinted against each of the 16 cortico-genetic correlation maps (Rg) from **Fig. 1**, against each of the 16 - log_10_ transformed cortico-genetic p-value maps (**Suppl. Fig. 3**) and against each of 16 vectors that concatenated thickness and area surface vectors, respectively (8 Rg maps concatenating area and thickness, 8 -log_10_(P) maps concatenating area and thickness). In sum, this yielded 48 Spearman correlation estimates (‘*fingerprinting strength’*) per subject. To assess the predictive utility of the fingerprinting strengths, we used those 48 correlations as features in machine learning based prediction of normative estimates of general psychopathology (pF) and general cognition (gF), respectively. PF and gF were obtained from a PCA following previous protocols^30^ from the full PNC sample (9490 individuals) and the respective scores extracted for those individuals with imaging data available. Dependencies with age were removed using locally weighted regression to account for non-linear effects. Machine learning was performed in a 10-fold cross-validation framework using *slm* from the *care* package^34^ in R statistics and normative estimates of pF and gF were predicted. Significance of the predictions was assessed using permutation testing, repeating 10,000 runs of a full 10-fold cross-validation loop using a different random permutation of the response variable in each run. Feature weights were assessed using *CAR scores* and translated to -log_10_ transformed p-values^34^. Finally, to assess environmental impact on cortico-genetic fingerprinting strength, we computed parental education as the mean of maternal and paternal education, and a principal component analysis across various trauma questions (**Suppl. Fig. 7**) yielding a general trauma score (the first factor). For each cortico-genetic map we tested for linear associations between individual fingerprinting strength and parental education accounting for age, sex and gF. Likewise, for each map we tested for linear associations with trauma, accounting for age, sex and pF (the pF was recomputed for this analysis to exclude trauma items). Significance of the linear associations was assessed using permutation testing, permuting the fingerprinting strength 10,000 times and each time recomputing the models on the permuted data. In addition, resulting P-values were corrected for multiple comparison using Bonferroni correction (p=.003125, 16 tests).

### Data availability

The data incorporated in this work are available as part of the publicly available UK Biobank (https://www.ukbiobank.ac.uk/) and Philadelphia Neurodevelopmental Cohort (PNC, https://www.ncbi.nlm.nih.gov/projects/gap/cgi-bin/study.cgi?study_id=phs000607.v2.p2).

### Code availability

Scripts are available upon request from the first author.

#### Supplementary figures

**Suppl. Fig.1:**
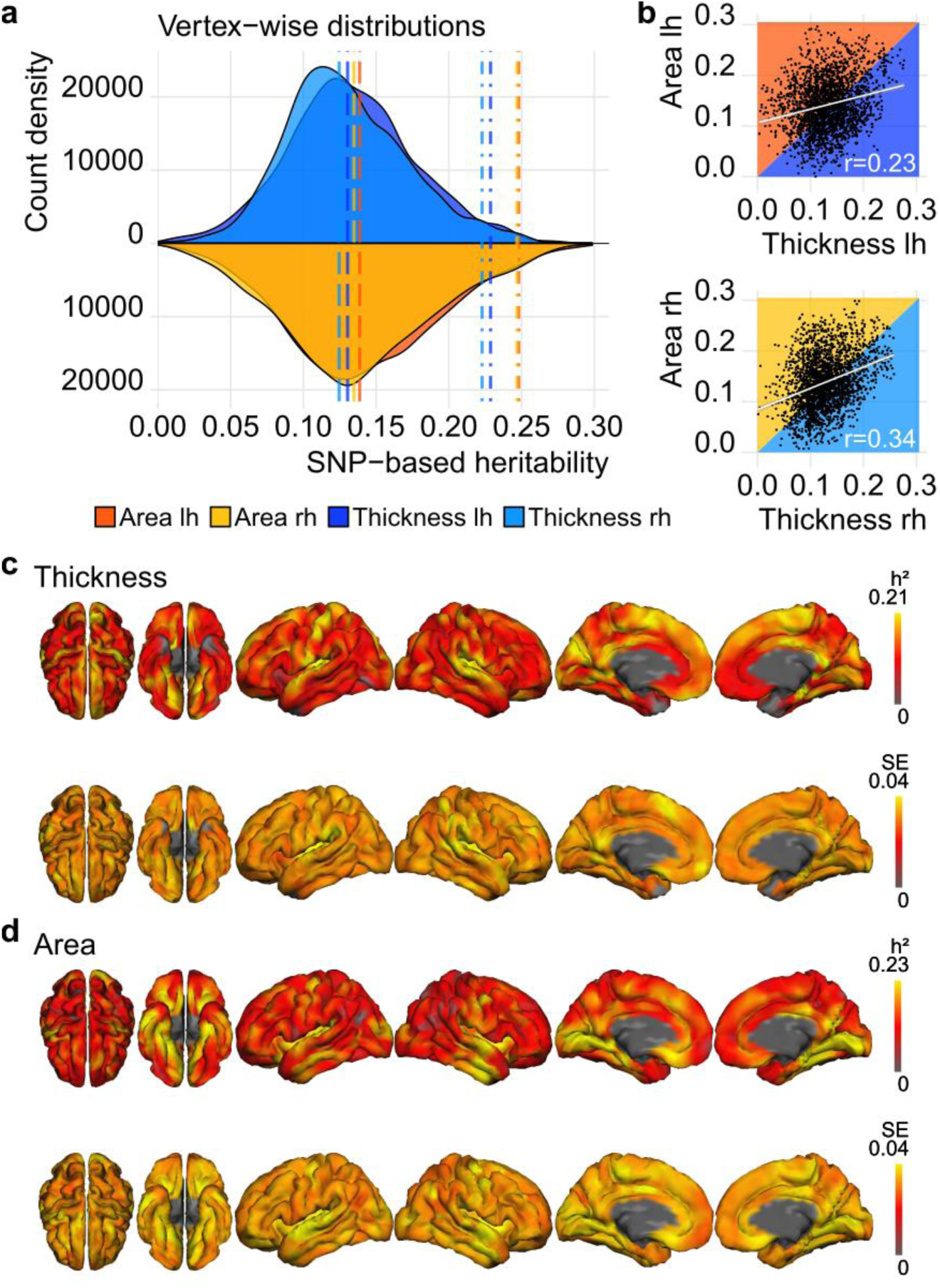
SNP-based heritability of cortical thickness and area confirms earlier reports from twin studies on heritability of cortical morphology. (a) Vertex-wise distribution of heritability estimates per hemisphere and cortical measure. Area was significantly more heritable than thickness (t=9.2, p<2e-16). To visualize this effect, the long-dashed lines indicate 50% quantiles of the distributions and the dot-dashed lines indicate the 97.5% quantiles. (b) Association of thickness and area heritability maps, per hemisphere. For both hemispheres concatenated, the association was r=0.28, p_perm_=.004 (c) Cortical maps for heritability of thickness (upper row) and corresponding standard error (lower row). (d) Same maps as (c), but for cortical area.

**Suppl. Fig.2:**
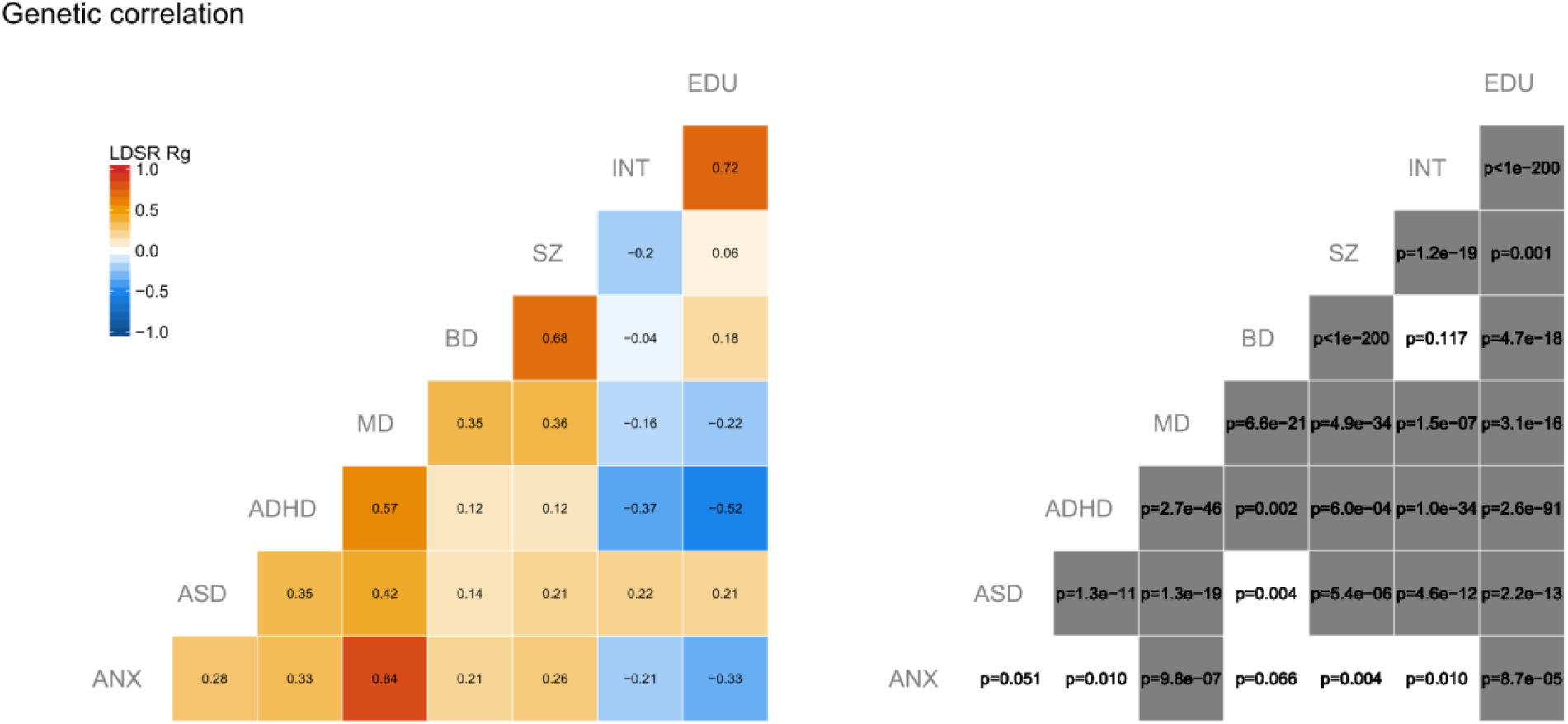
Genetic correlation matrix. Genetic correlation of summary statistics from different phenotypes using *LD-Score regression*^19,33^. The left plot shows the genetic correlations (Rg) and the right plot depicts the corresponding p-values. Significant associations following Bonferroni correction for the number of tests (28) are marked in grey.

**Suppl. Fig.3:**
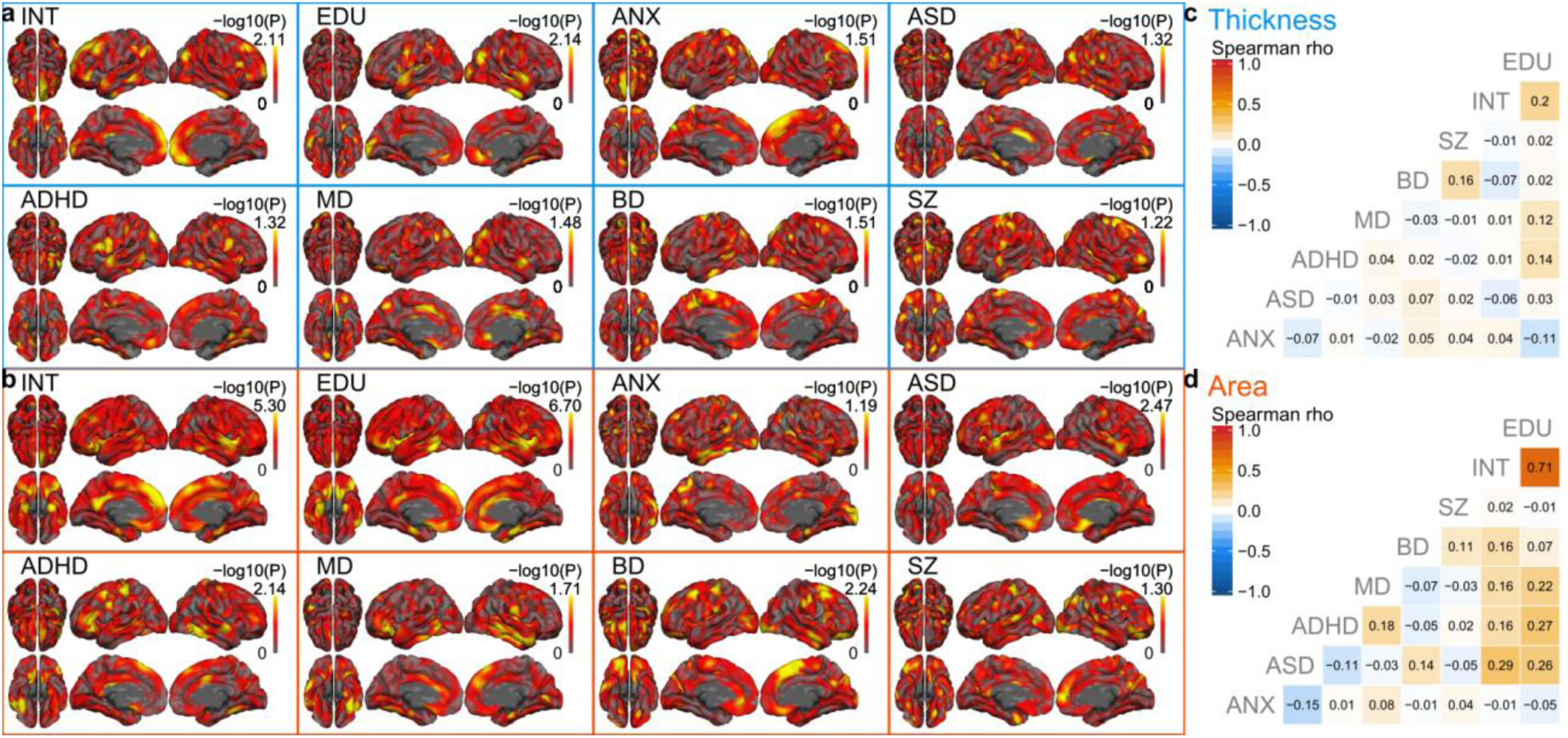
-log_10_ transformed p-values of cortico-genetic maps for cognition and psychiatric disorders. Corresponding with **Fig. 1** which displays the genetic correlations (Rg), the figures display the -log_10_ transformed p-values from vertex-wise *LD-score regression*^19,33^ for (a) thickness and (b) area. (c-d) Pairwise spearman correlations of each -log_10_ transformed p-value map.

**Suppl. Fig.4:**
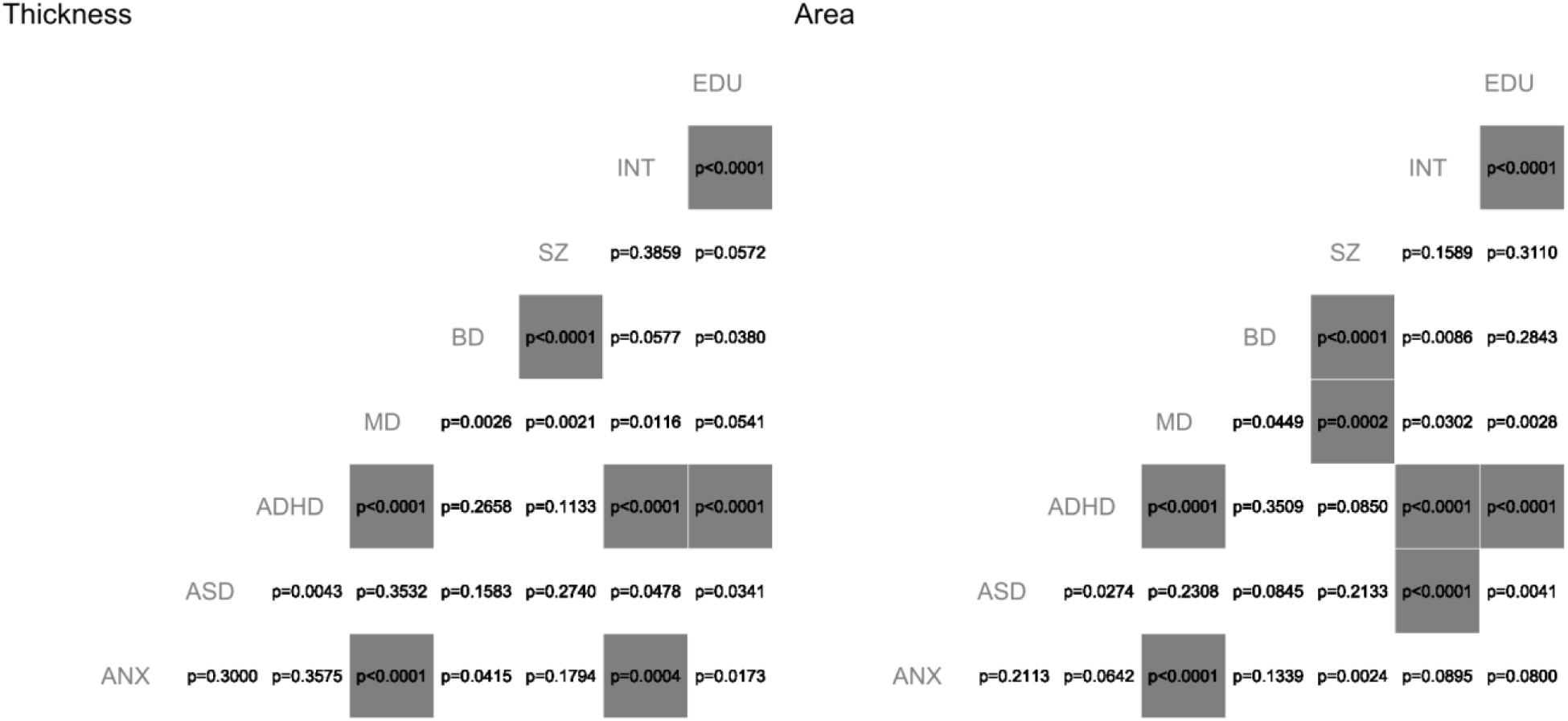
Permutation-based p-values corresponding to Fig. 1c (left) and 1d (right). Cortico-genetic maps were permuted using spin-rotation to derive a permutation-based p-value^32^. Significant associations following Bonferroni correction for the number of tests (28) are marked in grey.

**Suppl. Fig.5:**
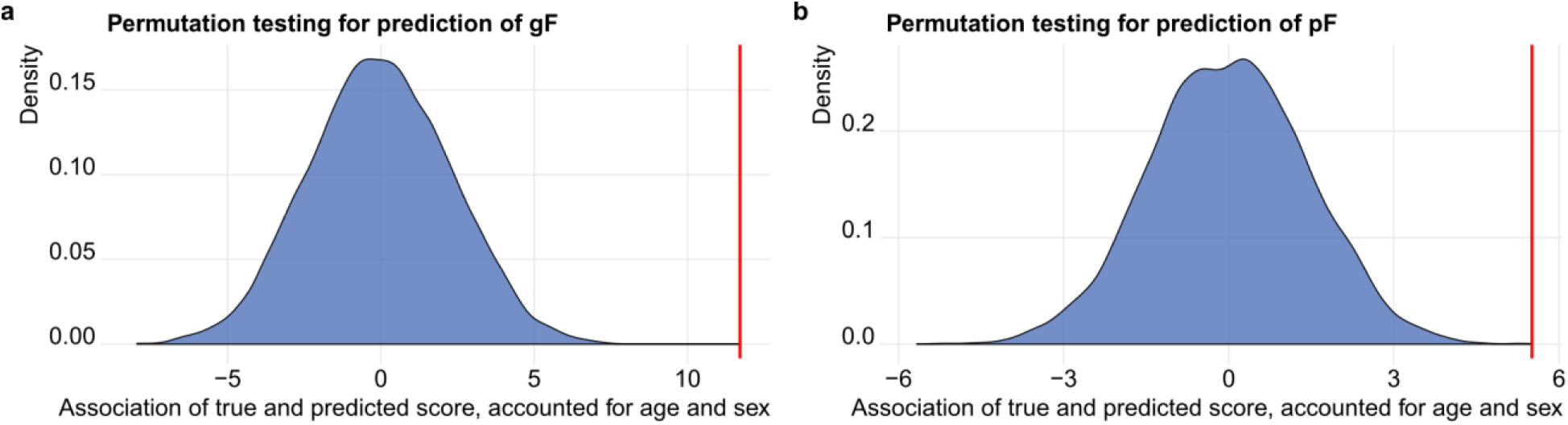
Permutation tests of the machine learning analyses confirms significant predictions of normative gF and normative pF. The red lines indicate the association between true and predicted scores (t-statistic), accounted for age and sex. The density plots depict the distribution of similar t-statistic obtained from 10,000 permutations per trait, with none of the permutation-based statistics exceeding the true value. (a) Significant prediction of normative gF (p_perm_<.0001). (b) Significant prediction of normative pF (p_perm_<.0001).

**Suppl. Fig.6:**
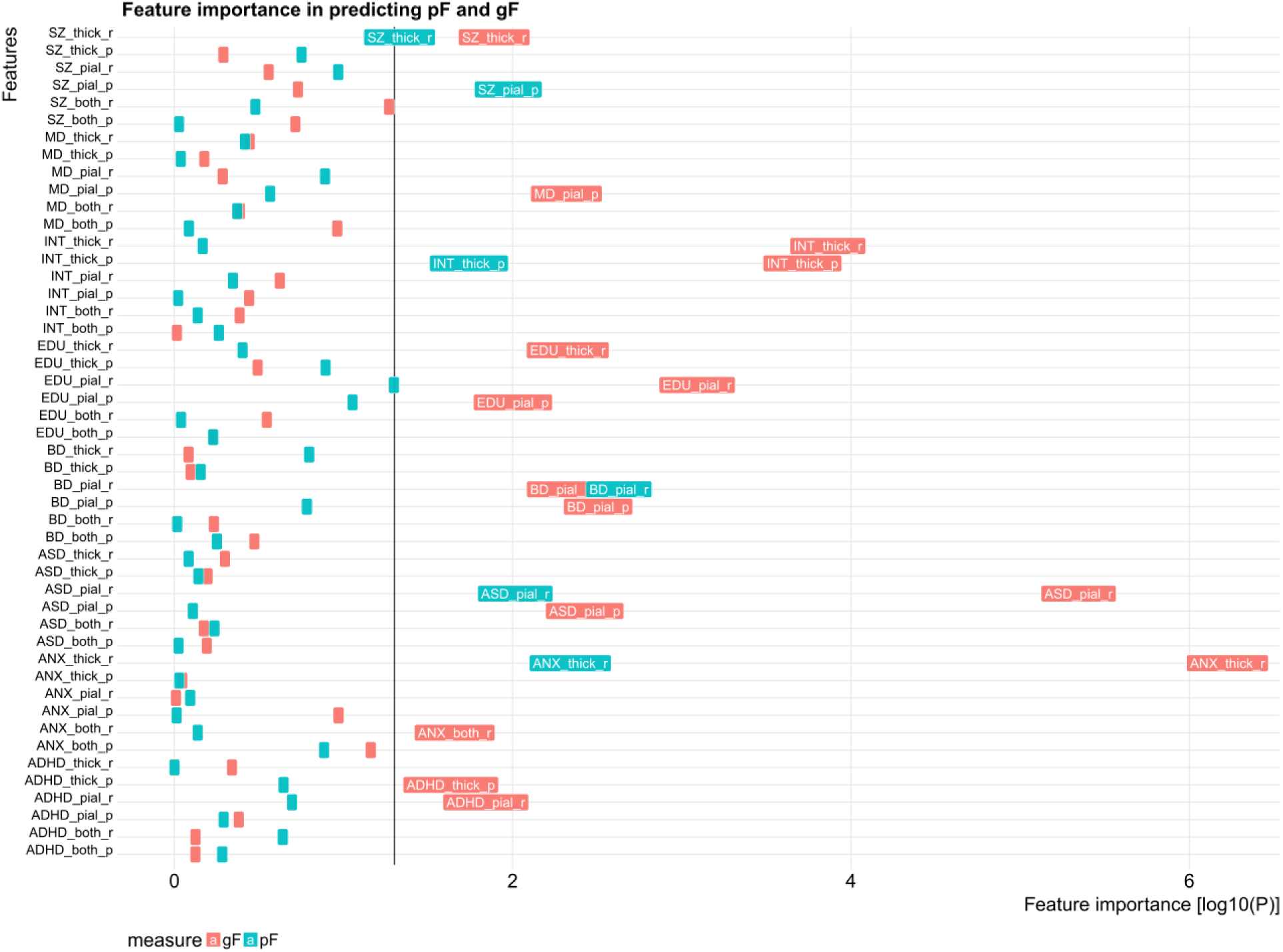
Feature importance. The -log_10_ transformed p-values of the CAR scores^34^ are displayed for all features. For visualization purpose only the most important features (p<.05) are labeled with text. Colors indicate the prediction model for normative gF (red) and normative pF (cyan).

**Suppl. Fig.7:**
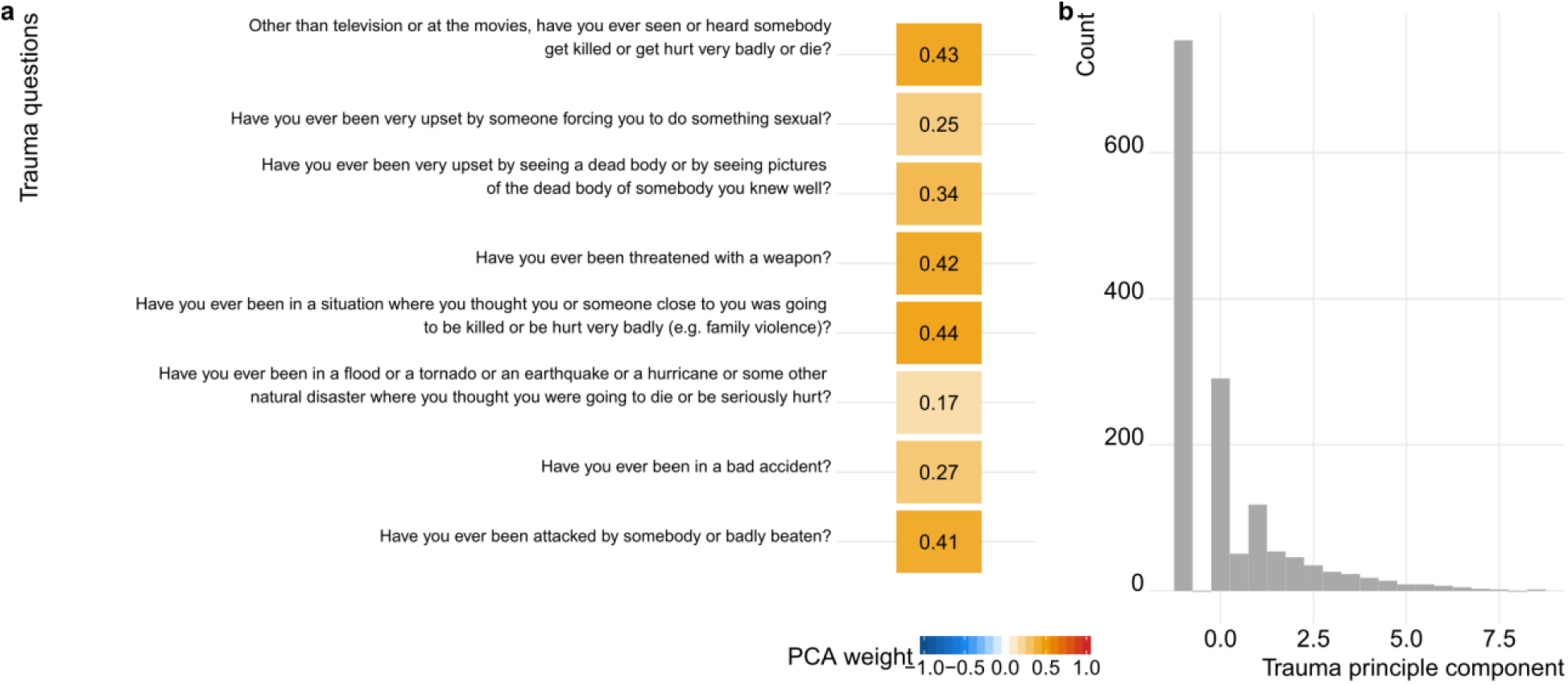
Details on the trauma principle component. (a) Trauma question and corresponding item weight in the PCA (b) Distribution of the trauma principle component across individuals.

**Suppl. Fig.8:**
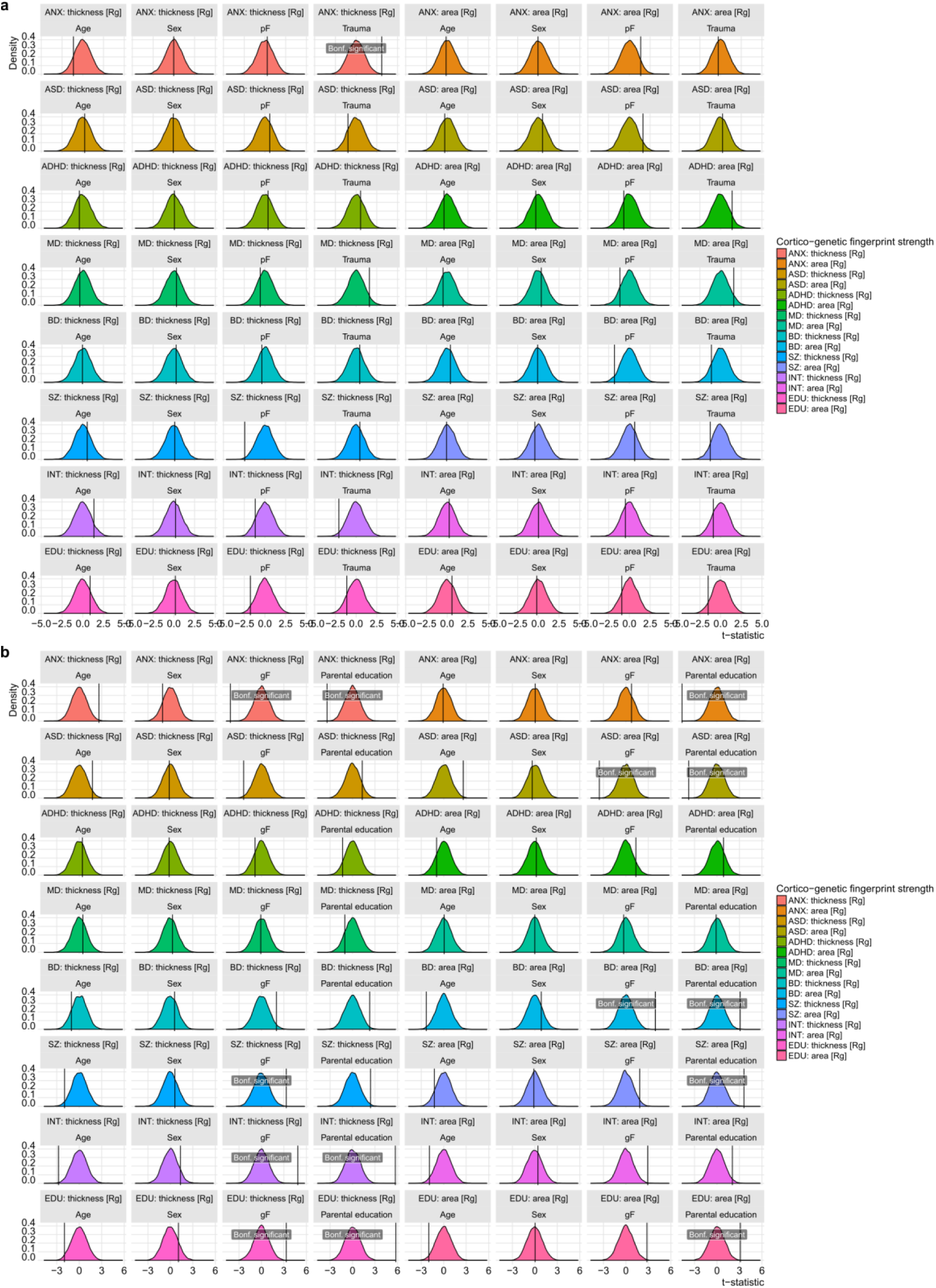
Permutation test results corresponding to Fig. 3. The cortico-genetic fingerprinting strength was permuted 10,000 times, each time calculating (a) the effect of age, sex, pF (excl. trauma) and trauma, and (b) the effect of age, sex, gF and parental education. The density plots illustrate the distribution of respective t-statistics. The vertical line indicates the true association t-statistic.

## References

1. Savage, J.E., et al. Nature genetics 50, 912–919 (2018).

2. Lee, J.J., et al. Nature genetics 50, 1112–1121 (2018).

3. Otowa, T., et al. Molecular psychiatry 21, 1391–1399 (2016).

4. Grove, J., et al. bioRxiv [https://doi.org/10.1101/224774] (2017).

5. Demontis, D., et al. Nature genetics (2018).

6. Wray, N.R., et al. Nature genetics 50, 668–681 (2018).

7. Stahl, E., et al. bioRxiv [https://doi.org/10.1101/173062] (2018).

8. Schizophrenia Working Group of the PGC, et al. Nature 511, 421 (2014).

9. Rimol, L.M., et al. Biological psychiatry 67, 493–499 (2010).

10. Eyler, L.T., et al. Twin Research and Human Genetics 15, 304–314 (2012).

11. Peper, J.S., Brouwer, R.M., Boomsma, D.I., Kahn, R.S. & Hulshoff Pol, H.E. Hum Brain Mapp 28, 464–473 (2007).

12. Panizzon, M.S., et al. Cerebral Cortex 19, 2728–2735 (2009).

13. Winkler, A.M., et al. Neuroimage 53, 1135–1146 (2010).

14. Sudlow, C., et al. PLoS Medicine 12, e1001779 (2015).

15. Satterthwaite, T.D., et al. Neuroimage 124, 1115–1119 (2016).

16. Fischl, B., et al. Neuron 33, 341–355 (2002).

17. Purcell, S., et al. American Journal of Human Genetics 81, 559–575 (2007).

18. Bycroft, C., et al. bioRxiv [https://doi.org/10.1101/166298] (2017).

19. Bulik-Sullivan, B.K., et al. Nature genetics 47, 291–295 (2015).

20. Grasby, K.L., et al. bioRxiv [https://doi.org/10.1101/399402] (2018).

21. Insel, T.R. Nature 468, 187–193 (2010).

22. Lee, F.S., et al. Science 346, 547–549 (2014).

23. Dahl, R.E., Allen, N.B., Wilbrecht, L. & Suleiman, A.B. Nature 554, 441–450 (2018).

24. Hastie, T. Generalized Additive Models. R package v 1.16 (2018).

25. Finn, E.S., et al. Nature neuroscience 18, 1664–1671 (2015).

26. Kaufmann, T., et al. Nature neuroscience 20, 513 (2017).

27. Miranda-Dominguez, O., et al. PloS one 9, e111048 (2014).

28. Schnack, H.G., et al. Cerebral Cortex 25, 1608–1617 (2015).

29. van Erp, T.G.M., et al. Biological psychiatry 84, 644–654 (2018).

30. Alnaes, D., et al. Jama Psychiat 75, 287–295 (2018).

## Supplementary references

1. Savage, J.E., et al. Nature genetics 50, 912–919 (2018).

2. Lee, J.J., et al. Nature genetics 50, 1112–1121 (2018).

3. Otowa, T., et al. Molecular psychiatry 21, 1391–1399 (2016).

4. Grove, J., et al. bioRxiv [https://doi.org/10.1101/224774] (2017).

5. Demontis, D., et al. Nature genetics (2018).

6. Wray, N.R., et al. Nature genetics 50, 668–681 (2018).

7. Stahl, E., et al. bioRxiv [https://doi.org/10.1101/173062] (2018).

8. Schizophrenia Working Group of the PGC, et al. Nature 511, 421 (2014).

14. Sudlow, C., et al. PLoS Medicine 12, e1001779 (2015).

15. Satterthwaite, T.D., et al. Neuroimage 124, 1115–1119 (2016).

16. Fischl, B., et al. Neuron 33, 341–355 (2002).

17. Purcell, S., et al. American Journal of Human Genetics 81, 559–575 (2007).

18. Bycroft, C., et al. bioRxiv [https://doi.org/10.1101/166298] (2017).

19. Bulik-Sullivan, B.K., et al. Nature genetics 47, 291–295 (2015).

24. Hastie, T. Generalized Additive Models. R package v 1.16 (2018).

25. Finn, E.S., et al. Nature neuroscience 18, 1664–1671 (2015).

26. Kaufmann, T., et al. Nature neuroscience 20,513 (2017).

27. Miranda-Dominguez, O., et al. PloS one 9, e111048 (2014).

30. Alnaes, D., et al. Jama Psychiat 75, 287–295 (2018).

31. Satterthwaite, T.D., et al. Neuroimage 86, 544–553 (2014).

32. Alexander-Bloch, A., et al. Neuroimage (2018).

33. Bulik-Sullivan, B., et al. Nature genetics 47, 1236–1241 (2015).

34. Zuber, V. & Strimmer, K. Care. R package v 1.1.10 (2017).

